# Computational Design of Soluble CCR8 Analogues with Preserved Antibody Binding

**DOI:** 10.1101/2025.08.18.670068

**Authors:** Trang Nguyen, Songming Liu, Yifan Li, Longfei Cong, Roger Shek, Tek Hyung Lee, Li Yi, Per Greisen

**Affiliations:** BioMap Research, Palo Alto, California, USA; BioMap Research, Beijing, China

**Keywords:** membrane protein engineering, GPCR solubilization, computational protein design, CCR8, antibody binding, surface plasmon resonance

## Abstract

G protein-coupled receptors (GPCRs) represent the largest class of drug targets, yet their membrane-embedded nature poses significant challenges for structural studies and therapeutic development. Here, we report the successful computational design and experimental validation of soluble CCR8 analogues that maintain native antibody binding properties. Using an integrated pipeline combining ProteinMPNN-sol sequence design, and structure-based filtering, we generated 13 CCR8 analogues from 272 initial designs across three N-terminal truncation strategies. Experimental validation confirmed 62% success rate (8/13 designs) with protein yields of 1.19-73.72 mg/L in aqueous buffer, representing a significant improvement over traditional membrane protein production method. Surface plasmon resonance analysis demonstrated that all analogues retained mAb1 binding with dissociation constants ranging from 77-857 nM, comparable to wild-type CCR8 (K_D_ = 190 nM). Despite extensive sequence divergence (10-13% identity with wild-type CCR8), structural integrity was preserved as evidenced by binding affinity maintenance and computational structural validation. This work demonstrates the feasibility of computationally designing functional soluble analogues of challenging membrane proteins, with implications for accelerating drug discovery, antibody development, and structural biology studies. Our approach addresses critical limitations in membrane protein accessibility while preserving native epitope presentation, opening new avenues for therapeutic target characterization and binder discovery.

## 1. Introduction

G protein-coupled receptors (GPCRs) constitute the largest family of cell surface receptors, mediating critical physiological processes and representing approximately 35% of all marketed drugs [14]. Despite their therapeutic importance, membrane proteins including GPCRs present formidable challenges for structural and functional studies due to their inherent hydrophobicity and requirement for lipid bilayer environments [15, 16]. Recent breakthrough advances in computational protein design have opened unprecedented opportunities for engineering soluble analogues of membrane proteins while preserving their native functional properties [17].

### 1.1 Challenges in Membrane Protein Research

The primary obstacles in membrane protein research stem from their unique biophysical properties. Their inherent hydrophobicity necessitates extraction from lipid bilayers using detergents or other membrane mimetics, often leading to protein instability and aggregation [2, 3]. Low expression levels in heterologous systems, coupled with difficulties in maintaining native conformations outside membrane environments, severely limit the quantities of purified, functional protein available for downstream applications including structural determination, biochemical assays, and antibody generation [2, 4].

These challenges extend significantly to therapeutic antibody development against membrane proteins. The limited accessibility of immunogenic regions embedded within lipid bilayers, combined with the small size of extracellular loops and conformational sensitivity of epitopes, often results in low immunogenicity and difficulties in eliciting antibodies that recognize native, functional protein states [5, 6]. Traditional immunization strategies using purified, denatured proteins or synthetic peptides frequently fail to produce antibodies binding conformational epitopes essential for biological activity [7]. Furthermore, immunization with full-length membrane protein constructs presents additional hurdles including low surface expression levels, potential cellular toxicity, and complexity in maintaining structural integrity outside native membrane environments [5, 7].

### 1.2 Revolutionary Advances in Computational Design

Recent computational breakthroughs have fundamentally transformed membrane protein engineering capabilities. The landmark 2024 work by Goverde et al. demonstrated the first successful computational design of soluble analogues of integral membrane proteins using deep learning approaches [17]. Their AF2seq-ProteinMPNN pipeline achieved remarkable success rates of 77% for claudin-like folds, 87% for rhomboid protease folds, and 64% for GPCR-like folds, while preserving native functionality as evidenced by crystal structures showing design accuracy within 0.73-1.05 Å RMSD [17].

The integration of AlphaFold2-based sequence design (AF2seq) with ProteinMPNN-sol represents a paradigm shift in protein engineering [18, 19]. AF2seq inverts the AlphaFold2 network using gradient descent optimization with composite loss functions, generating highly novel sequences through iterative refinement [17]. ProteinMPNN-sol, specifically trained on soluble proteins, addresses critical limitations of standard ProteinMPNN when applied to membrane protein topologies [19]. Advanced solubility prediction tools including SoluProt, provide robust computational assessment of design success probability [21, 22, 23].

### 1.3 CCR8 as a Critical Therapeutic Target

Among the GPCR family, CCR8 (C-C chemokine receptor 8) has emerged as a particularly important therapeutic target due to its central role in immune regulation and disease progression. CCR8 is predominantly expressed on regulatory T cells (Tregs), which maintain immune tolerance and suppress anti-tumor immune responses within the tumor microenvironment [10, 11]. Dysregulation of CCR8 and its primary ligand CCL1 has been implicated in multiple cancer types including breast invasive carcinoma, colon adenocarcinoma, and head and neck squamous cell carcinoma, where it promotes immunosuppressive environments [10, 12].

Targeting CCR8 represents a promising cancer immunotherapy strategy aimed at disrupting Treg recruitment and enhancing anti-tumor immunity [11, 13]. Beyond oncology applications, CCR8 modulators are being explored for treating inflammatory and autoimmune diseases [13]. However, the membrane-embedded nature of CCR8 has limited structural studies and antibody development efforts, highlighting the critical need for soluble, functional analogues.

### 1.4 Study Objectives and Innovation

This report demonstrates the successful computational design and experimental validation of soluble CCR8 analogues that retain native antibody binding properties. By integrating state-of-the-art computational tools like ProteinMPNN-sol and structure-based filtering with comprehensive experimental validation through surface plasmon resonance and flow cytometry, we address fundamental limitations in membrane protein accessibility while preserving critical epitope presentation. This work represents a significant advancement in making CCR8 and similar membrane proteins more tractable for biochemical studies, antibody development, and therapeutic applications.

## 2. Materials and Methods

### 2.1 Computational Design Pipeline

We developed an integrated computational pipeline combining ProteinMPNN-sol and structure-based filtering criteria, building on recent advances in membrane protein solubilization [17, 19]. The pipeline systematically transforms membrane-embedded topologies into soluble analogues while preserving functional epitopes.

#### 2.1.1 Target Structure Preparation

The CCR8–mAb1 complex (PDB ID: 8TLM) was obtained from the Protein Data Bank. Interface residues involved in antibody binding were identified and fixed throughout the design process to preserve functional interactions. Specifically, CCR8 residues were selected if their backbone atoms were within 7 Å and side-chain atoms within 6 Å of the antibody, following established protocols for epitope preservation [24]. All candidate residues were manually curated to retain functionally important sites while excluding sterically unreasonable positions.

ProteinGenerator was employed to reconstruct missing N-terminal residues (positions 1-32) in the crystal structure [25]. To evaluate N-terminal length impact on design success, three truncation strategies were implemented: (i) full-length N-terminus (Truncation Strategy 1(TS1): WT 1-314), (ii) truncation at V9_TTV (Truncation Strategy 2(TS2): WT 10-314), and (iii) truncation at D26_AE (Truncation Strategy 3(TS3): WT 27-314).

#### 2.1.2 ProteinMPNN-sol Sequence Redesign

For each trajectory, the sequence with lowest final loss was selected and re-folded using xTrimoMonomer(unpublished) structure prediction. Predicted structures were evaluated using stringent quality criteria: TM-score > 0.75 and Cα RMSD of fixed residues < 4 Å. High-confidence models were used as structural templates for sequence redesign with ProteinMPNN-sol v_48_020, constraining fixed interface residues throughout the process [19]. For each structure, 16 new sequences were generated using sampling temperatures of 0.05, 0.1, 0.2, and 0.3 to balance sequence diversity with structural compatibility.

#### 2.1.4 Structure and Solubility Filtering

All redesigned sequences underwent comprehensive evaluation for structural quality and solubility. xTrimoMonomer was used for template-free structure prediction. The following stringent filtering criteria were applied:

1. No more than four consecutive repeats of the same amino acid residue, except for glutamate (up to five consecutive allowed), with no restrictions applied to cysteine residues
2. TM-score > 0.8 between designed structure and target topology
3. Mean pLDDT > 80
4. Cα RMSD of fixed residues < 3 Å relative to target structure

#### 2.1.5 Sequence Identity Calculation

To assess the diversity among designed sequences, we computed the pairwise sequence identity between all experimentally validated designs and the corresponding wild-type sequence. All sequences were aligned to the wild-type sequence based on residue positions for position-wise comparison.

For TS2, the wild-type reference was the CCR8 sequence from residue 10 to 314, while TS2 designs covered positions 27 to 314. For each pair of sequences with equal length, sequence identity was defined as the percentage of positions with identical amino acids:

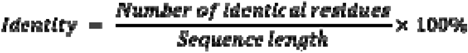

For sequence pairs of different lengths (e.g., designs from TS2 vs. TS3), the shorter sequence was padded to match the length of the longer one before computing identity. Only exact amino acid matches were considered; biochemical similarities were not included in the identity calculation.

### 2.2 Prescreening for binding of soluble CCR8 analogues with mAb1 via Yeast Surface Display

13 CCR8 soluble analogues that passed the computational filtering pipeline were expressed and purified in *E. coli* for prescreening of mAb1 binding using yeast surface display. Constructs with C-terminal His tag, were cloned into OverExpress C43(DE3) Chem Comp Cells (Biosearch Technologies) and grown in 4mL Luria-Broth at 37°C. Constructs with C-terminal His tag, were cloned into OverExpress C43(DE3) Chem Comp Cells (Biosearch Technologies) and grown in 4 mL Luria-Broth at 37°C. Cells were induced with 1 mM IPTG at OD_600 ∼_1, then expressed for 4 hours at 37 ° C prior to harvesting. Lysed cells were purified with Ni magnetic beads and eluted with 200mM imidazole. The purified proteins were subsequently buffer exchanged into PBS. Binding of CCR8 variants to mAb1 (displayed in scFv format on yeast) was assessed by flow cytometry before selecting candidates for large-scale production and surface plasmon resonance (SPR) characterization. Biotinylated Human CCR8 Protein, His,Flag&Avitag in detergent was purchased from Acro Biosciences (Cat. # CC8-H82D3).

### 2.3 Protein Production

Eight CCR8 soluble variants that passed pre-screening were scaled up and expressed by Biointron Biological Inc. Constructs were cloned into One Shot™ BL21 (DE3) Chemically Competent *E. coli* (Invitrogen) and grown in 1 L of Luria-Broth at 37°C. Cells were induced with 1 mM IPTG at OD_600 ∼_1, then expressed for 4 hours at 37 ° C prior to harvesting. Cells were lysed, and proteins were purified using HisTrap affinity chromatography followed by size-exclusion chromatography (SEC) and stored in PBS. For SPR study, mAb1 in full length human IgG1 format was produced in CHO cells, purified by one step Protein A chromatography and stored in PBS. Wild-type CCR8 used as an assay control was purchased from Acro Biosystems (Cat. #CC8-H52D6).

Protein identity and purity were characterized using SDS-PAGE analysis under reducing and non-reducing conditions and SEC-HPLC analysis.

### 2.4 Surface Plasmon Resonance Validation

Binding affinity measurements were performed using a Biacore 8K instrument (Cytiva) by Acro Biosystems. A Series S Sensor Chip Protein A (Cytiva, Cat. No. 29127556, Lot. No. 10359311) was utilized for mAb1 capture, following established SPR protocols for membrane protein characterization [26, 27].

The running buffer consisted of 1×HEPES (10 mM HEPES, 150 mM NaCl, 3 mM EDTA) with 0.005% Tween-20 at pH 7.4. For wild-type CCR8 stabilization, 0.05% DDM and 0.01% CHS were added to the running buffer. Regeneration was performed using 10 mM Glycine-HCl, pH 1.5. mAb1 capture was achieved by flowing 5 μg/mL antibody in running buffer at 10 μL/min, targeting capture levels of approximately 400 RU. Association and dissociation phases were monitored at 30 μL/min flow rate with 90-second association and 210-second dissociation times.

## 3. Results

### 3.1 Computational Design Pipeline Success

Our integrated computational pipeline successfully generated 13 CCR8 analogues from three design strategies across 272 *in silico* designs (Table 1). The systematic filtering process demonstrated the effectiveness of ProteinMPNN-sol redesign and structure-based validation, consistent with recent breakthrough advances in membrane protein solubilization [17].

**Table 1.**
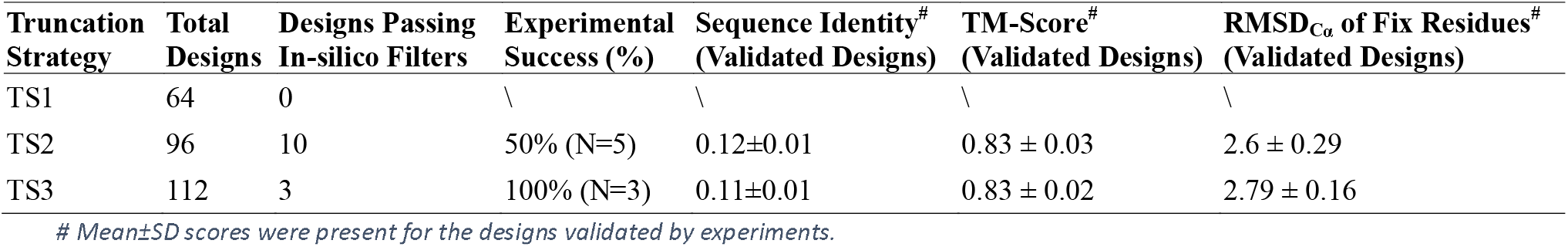
Computational Design Success Rates and Structural Validation Metrics.

The TS2 truncation strategy proved most successful, yielding 10 designs that passed our stringent filtering criteria with 5 experimentally validated as soluble (BM4 to BM8) (50% experimental success rate). The TS3 truncation approach generated 3 designs, all of which demonstrated experimental solubility (BM1 to BM3) (100% success rate). Notably, the TS1 strategy, which preserved the full-length N-terminus, failed to produce any designs meeting filtering criteria, suggesting that N-terminal truncation significantly impacts solubilization success, consistent with observations in other membrane protein engineering studies [37].

Sequence analysis revealed that successful designs maintained only 10-13% sequence identity with wild-type CCR8 while preserving structural integrity, as confirmed by TM-scores >0.8 and low RMSD values for fixed epitope residues (**Table 2**). This extensive sequence divergence while maintaining essential residues, demonstrates the robustness of our computational design approach.

**Table 2.** Production Yields and Purity Analysis of CCR8 Analogues and mAb1.

| Name | MW. (KDa) | Concentration (mg/mL) | Yield (mg/L) | SEC-HPLC Purity(%) |
| --- | --- | --- | --- | --- |
| BM1 | 32.30 | 0.31 | 1.19 | 94.68 |
| BM2 | 32.50 | 2.22 | 15.28 | 99.92 |
| BM3 | 32.60 | 1.86 | 10.18 | 99.16 |
| BM4 | 36.10 | 1.86 | 6.11 | 98.05 |
| BM5 | 35.60 | 1.52 | 22.69 | 92.32 |
| BM6 | 35.70 | 1.96 | 47.53 | 95.85 |
| BM7 | 35.40 | 3.45 | 73.72 | 92.3 |
| BM8 | 34.60 | 4.68 | 47.92 | 81.12 |
| mAb1 | 145.10 | 0.9 | 82.60 | 93.8 |

### 3.2 Structural Validation

Computational structural analysis using PyMOL alignment showed the designed analogues maintain the overall fold architecture (**Figure 6**). Predicted structures for BM3 and BM8 showed RMSD values of 4.08 Å and 3.66 Å respectively when aligned with crystal structure 8TLM, indicating successful preservation of core structural features despite extensive sequence modifications.

Sequence alignment and lipophilicity analysis revealed systematic transformation of the membrane protein architecture, with hydrophobic transmembrane regions converted to hydrophilic solvent-accessible regions to enable aqueous solubility while maintaining the overall structural framework (**Figure 7**). This fundamental topology conversion whereby replacing membrane-spanning hydrophobic segments with solvent-accessible hydrophilic alternatives, effectively inverts the protein’s environmental preferences from lipid bilayer to aqueous solution. Detailed sequence alignment analysis revealed systematic hydrophobic-to-hydrophilic substitutions across all designs while preserving critical epitope residues (Supplementary Figure 2). Critical epitope residues were preserved across all designs, validating the computational constraint strategy.

### 3.3 Prescreening for binding of soluble CCR8 Analogues with mAb1 by Yeast Surface Display

To assess solubility and retention of mAb1 binding in our computationally designed CCR8 variants, small-scale expression was performed in *E. coli*. Of the 13 designs that passed our filtering pipeline, eight were successfully expressed and purified in a single step using Ni magnetic beads. Binding was then evaluated using mAb1 (scFv format), displayed on the yeast surface, with CCR8 tested at two concentrations (25 nM and 250 nM). Several variants exhibited stronger binding at 250 nM compared to 25 nM, indicating a concentration-dependent increase in binding. Meanwhile, wild-type CCR8 in detergent showed robust binding at 50 nM to yeast-displayed mAb1, consistent with literature reports of sub-nanomolar affinity [30], confirming the sensitivity of our assay. Based on these results, the eight variants were selected for large-scale production and subsequent SPR characterization.

### 3.4 Large-scale Protein Production

The production of CCR8 analogues from Biointron Biological Inc. also confirmed 62% overall success rate (8/13 designs) for soluble membrane protein expression, indicating significant improvements over traditional approaches [7]. Protein yields ranged from 1.19-73.72 mg/L in aqueous PBS buffer, representing 10- to 100-fold improvement over typical detergent-solubilized membrane protein preparations which often yield <1 mg/L [36], with several analogues (BM5, BM6, BM7, BM8) achieving yields > 22 mg/L suitable for comprehensive biochemical characterization.

SEC-HPLC analysis demonstrated high purity levels across all analogues, with most showing >90% purity (**Table 2**). SDS-PAGE analysis confirmed expected molecular weights and absence of significant aggregation or degradation products (**Figure 3**). The successful production of multiple high-quality analogues validates the computational predictions and demonstrates the high practical effectiveness of our design approach.

**Figure 1.**
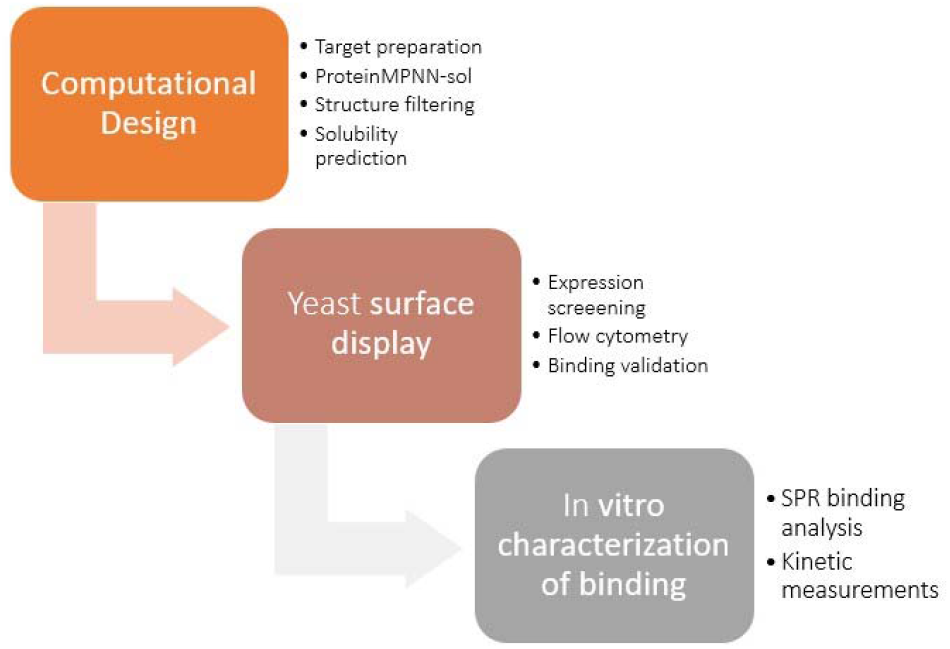
Integrated Pipeline for CCR8 Receptor Solubilization and Validation. The computational design workflow consists of three main stages: (1) **Computational Design** - Starting from the CCR8-mAb crystal structure (PDB: 8TLM), 16-18 critical epitope residues were fixed followed by ProteinMPNN-sol redesign. Stringent filtering criteria (TM-score >0.8, pLDDT >80) selected 13 final designs from 272 initial candidates across three N-terminal truncation strategies. (2) **Yeast Surface Display** - Selected 13 final designs underwent small-scale expression in E.coli and screened against surface displayed mAb1 (scFv format) using flow cytometry to validate structural epitope preservation. (3) **In Vitro Characterization** - Eight of 13 CCR8 analogues (BM1-BM8), which were confirmed soluble from small scale expression and bound with mAb1 by YSD, were produced at scale (1.19-73.72 mg/mL yields), characterized by SDS-PAGE and SEC-HPLC for purity, and quantitatively analyzed by Surface Plasmon Resonance (SPR) to determine mAb1 binding affinities (K_D_ = 77-857 nM). The pipeline achieved 62% experimental success rate (8/13 designs soluble) while maintaining antibody binding despite 10-13% sequence identity with wild-type CCR8, demonstrating effective epitope preservation and solubilization of a challenging membrane protein target.

**Figure 2.**
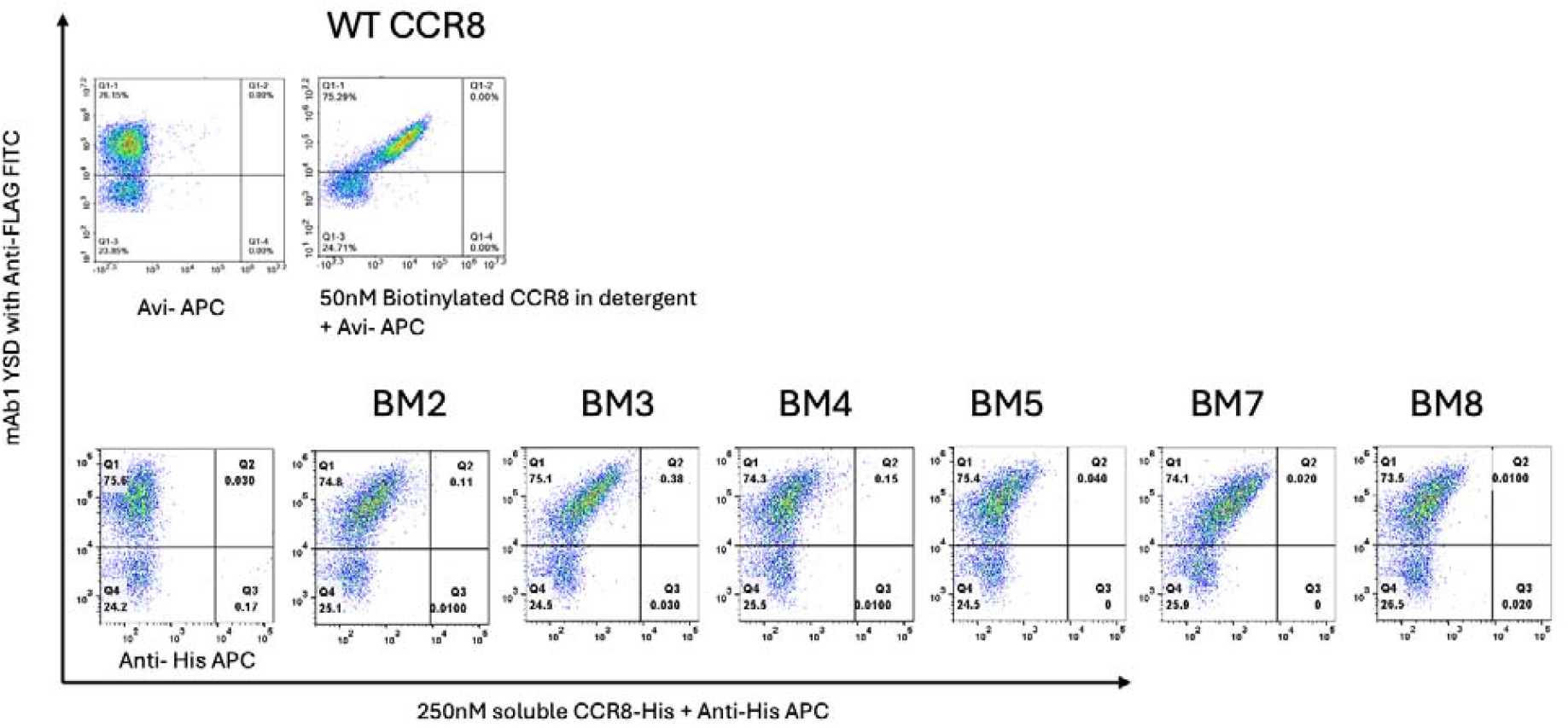
Binding Characterization of WT and soluble CCR8 Analogues with mAb1 by Flow Cytometry. Flow cytometry plot demonstrating the binding of 250 nM CCR8 Analogues (BM 2, 3, 4, 5, 7, 8) and 50 nM WT CCR8 to surface-displayed mAb1 (scFv form). The y-axis shows the display efficiency of mAb1 on the yeast surface, while the x-axis represents mAb1 binding to biotinylated WT and CCR8-His tag analogues detected via Avi and His tag Antibody, Alexa Fluor 647, respectively.

**Figure 3.**
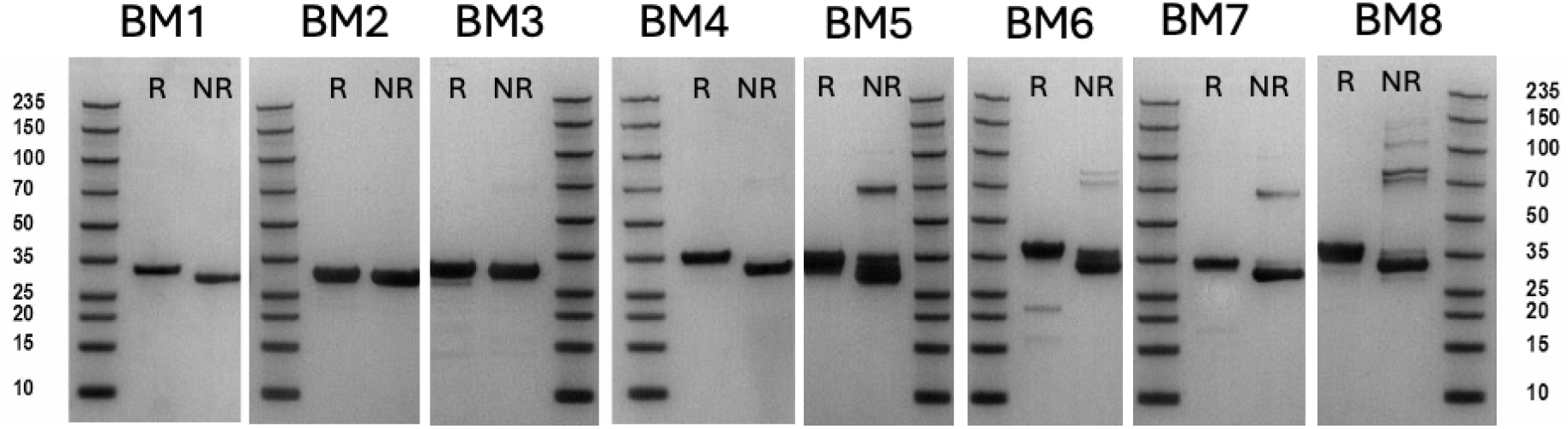
SDS-PAGE Analysis of Solubilized CCR8 Analogue Proteins. SDS-PAGE gel shows the purity and approximate molecular weights of the solubilized CCR8 analogue proteins used for SPR characterization.

### 3.5 Binding Validation by Surface Plasmon Resonance

Surface plasmon resonance analysis revealed that all eight CCR8 analogues retained mAb1 binding capability with dissociation constants ranging from 7.71×10□□ M to 8.57×10□□ M (77-857 nM) (**Table 3, Figure 4**). Notably, analogues BM2 and BM8 exhibited the highest binding affinity with K_D_ values of 77 nM and 88 nM, respectively, demonstrating comparable binding relative to wild-type CCR8 (K_D_ = 190 nM). The observed K_D_ value for WT-CCR8 in this study (190 nM) is higher than previously reported mAb1 affinity of 28.4 pM for cell-surface CCR8 [30]. This difference may be caused by using detergent-solubilized protein versus native membrane presentation, which can significantly impact binding kinetics and measured affinities [31]. Similar effects have been observed in other membrane protein SPR studies, where solubilized proteins show reduced apparent affinities compared to native membrane contexts.

**Table 3.** SPR Binding Affinity Data. Complete SPR kinetic parameters for all CCR8 analogues and wild-type CCR8 binding to mAb1, including KD values, kinetic constants, and quality metrics.

| Constructs | KD (M) | Rmax (RU) | Chi² (RU²) |
| --- | --- | --- | --- |
| BM1 | 2.67E-07 | 143.3 | 2.92 |
| BM2 | 7.71E-08 | 153.0 | 0.86 |
| BM3 | 1.54E-07 | 155.3 | 0.89 |
| BM4 | 8.57E-07 | 155.9 | 0.16 |
| BM5 | 6.29E-07 | 145.4 | 0.73 |
| BM6 | 5.52E-07 | 130.1 | 1.82 |
| BM7 | 2.52E-07 | 155.0 | 0.86 |
| BM8 | 8.79E-08 | 159.7 | 0.39 |
| WT-CCR8 | 1.90E-07 | 408.8 | 6.41 |

**Figure 4.**
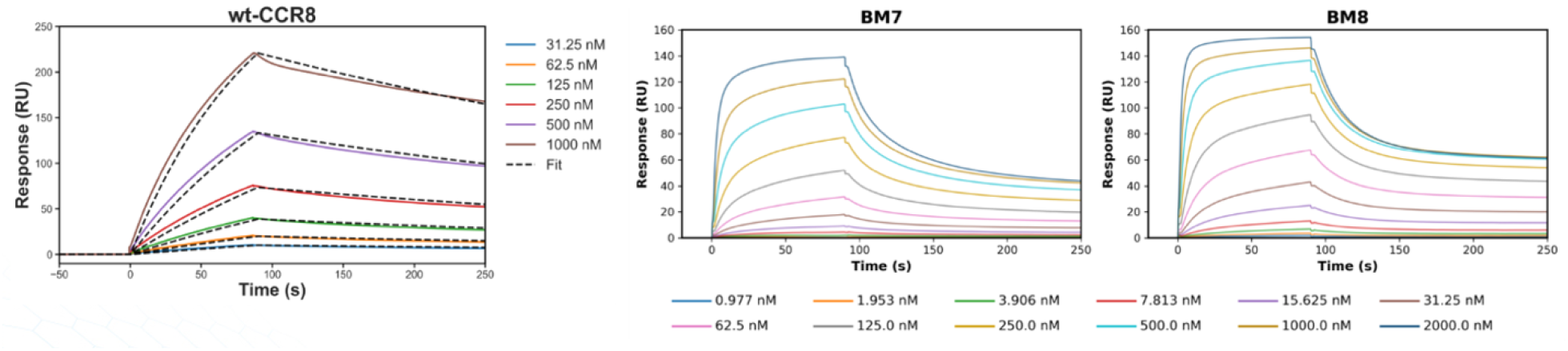
Surface Plasmon Resonance Binding Kinetics. Representative SPR sensorgrams showing real-time binding interactions between captured mAb1 and CCR8 analogues (left: wild-type CCR8, center: BM7, right: BM8). Concentration series demonstrate dose-dependent binding with clear association and dissociation phases. Wild-type CCR8 required detergent stabilization (DDM/CHS), while soluble analogues were analyzed in aqueous buffer, highlighting the practical advantages of the designed variants.

The binding affinity range spans approximately one order of magnitude, indicating that computational design successfully preserved core epitope interactions while allowing variation in binding strength (**Figure 5**). Analogue BM4 exhibited the weakest binding (857 nM), still within physiologically relevant ranges for antibody-antigen interactions. Chi-squared values for most measurements were <1.0 RU^2^, indicating excellent fit quality and reliable binding data.

**Figure 5.**
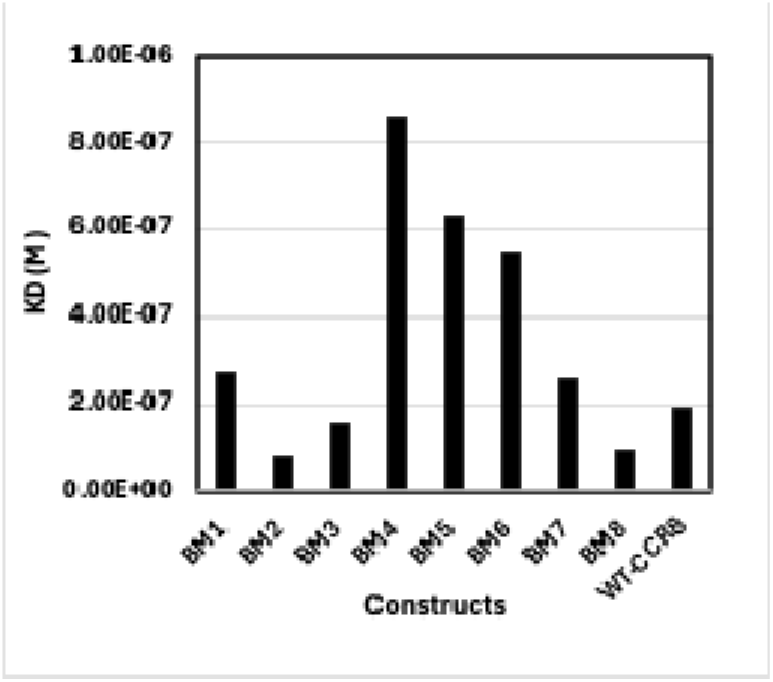
KD Values for CCR8 Analogue and Human CCR8. A bar graph illustrating the dissociation constants (KD) for the binding of mAb1 to eight computationally designed CCR8 analogues and the wild-type Human CCR8 protein, as determined by Surface Plasmon Resonance (SPR) using a steady-state affinity fit model. Lower KD values indicate tighter binding affinity.

### 3.6 Structure-Function Relationships

Analysis of binding affinity versus sequence conservation revealed important structure-function relationships (**Figure 6 and 7**). BM2 and BM8, showing the tightest binding affinities (77 and 88 nM), exhibit higher inter-analogue sequence identity (Figure 7B), exhibited greater compatibility with antibody recognition, likely due to their enhanced ability to stabilize the conformation of the recognized epitope while minimizing structural disruptions caused by solubilization. This suggests that their transition from membrane-bound to soluble forms preserved key polar and charged residues, maintaining an environment conducive to effective antibody engagement. Conversely, BM4 with the weakest binding (857 nM) may reflect suboptimal stabilization of the epitope region in the aqueous environment, where extensive sequence modifications could have disrupted the local conformational stability or introduced unfavorable interactions that compromise mAb1 recognition.

**Figure 6.**
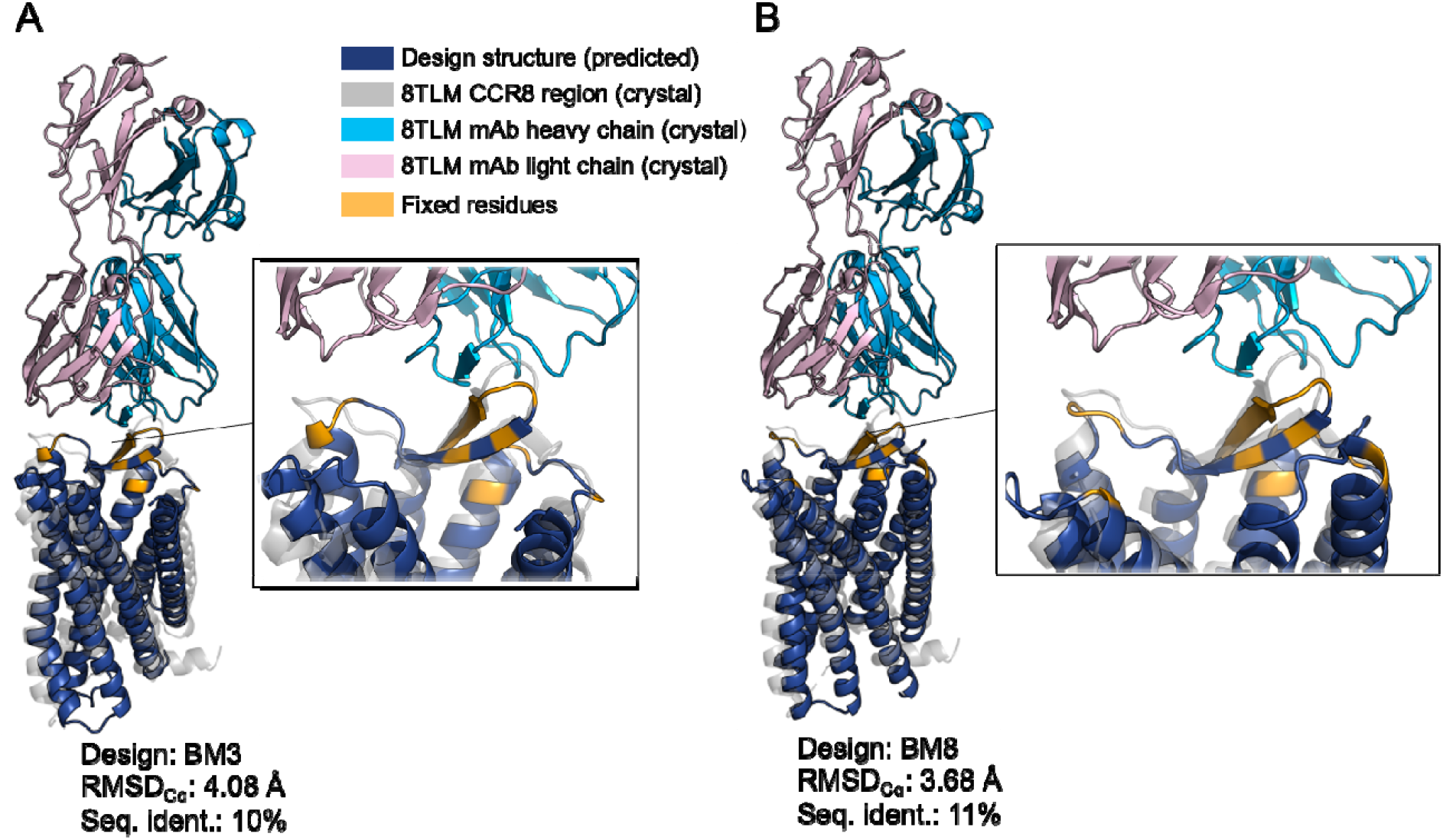
Structure Alignments of Designed Soluble CCR8 Analogues with Crystal Structure. Structure alignments comparing predicted BM3 (A) and BM8(B) with the crystal CCR8 structure (PDB ID: 8TLM) sourced from PDB. Although the soluble designs have a lower sequence identity to wild-type CCR8, their structural conformations are largely maintained. RMSD was calculated using PyMOL super command with cycles set to 0.

**Figure 7.**
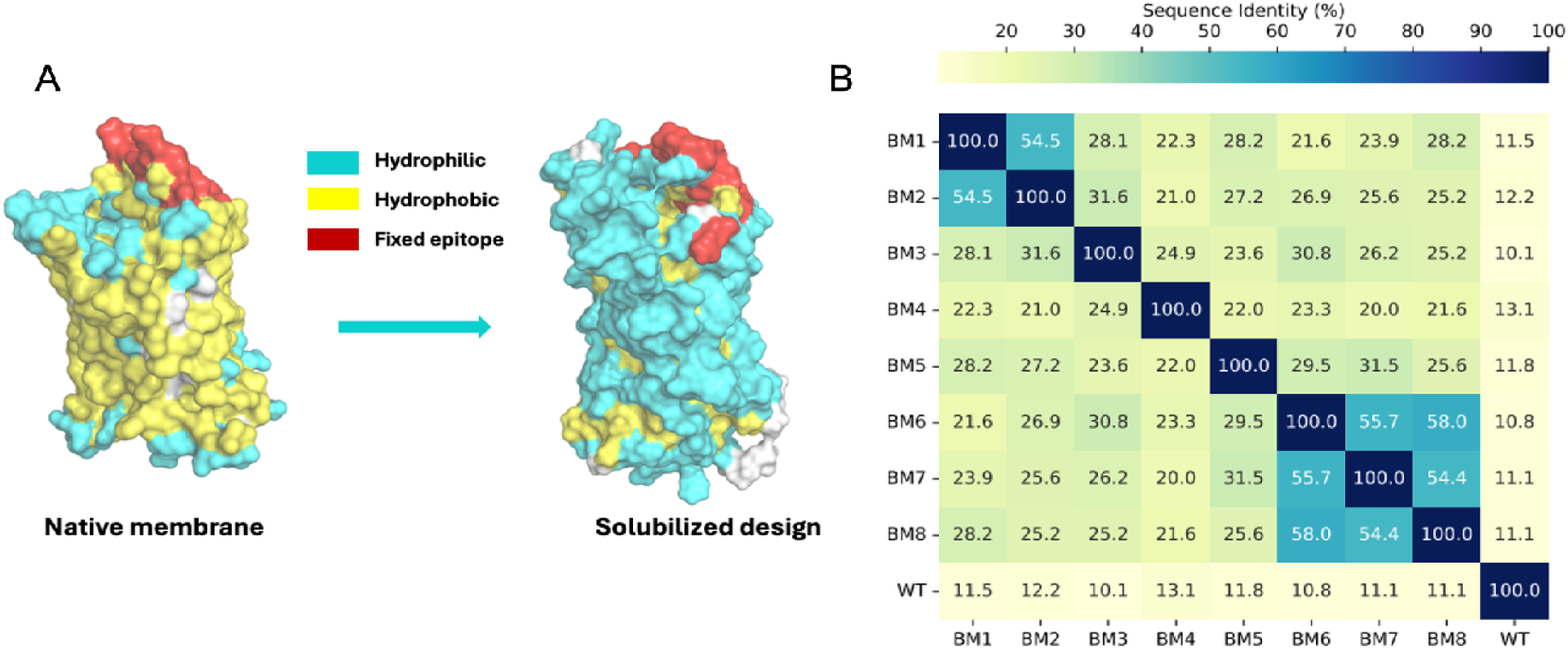
Comparison of Designed Soluble CCR8 Analogues and Wild-Type CCR8. (A) The example of amino acid change patterns. The hydrophobic surface residues in wild-type CCR8 were replaced by hydrophilic residues. (B) The sequence identity among soluble designs and wild-type CCR8.

## 4. Discussion

### 4.1 Breakthrough Achievement in Membrane Protein Engineering

This study demonstrates successful computational design of functional soluble CCR8 analogues, representing a significant advancement in membrane protein engineering. The 62% experimental success rate compares favorably with recent breakthrough studies, including the landmark Goverde et al. work that achieved 64% success for GPCR-like folds [17]. Our results validate the broader applicability of ProteinMPNN pipeline to challenging therapeutic targets while maintaining functional epitope presentation.

The preservation of antibody binding despite 10-13% sequence identity with wild-type CCR8 demonstrates remarkable robustness of the computational design approach (**Figure 6 and Supplemental Figure 2**). This extensive sequence divergence while maintaining function aligns with recent findings showing that membrane protein analogues can achieve native-like activities with minimal sequence conservation [17]. Our success validates computational epitope preservation strategies and suggests broad applicability to other membrane protein targets.

### 4.2 Methodological Advances and Technical Innovation

The systematic exploration of N-terminal truncation strategies provides valuable insights into design space optimization, with clear evidence that extensive truncation (TS3, TS2) significantly improves success rates compared to full-length constructs (TS1).

The stringent filtering criteria combining structural quality metrics (TM-score, pLDDT, RMSD) effectively enriched for experimentally successful designs. This multi-criteria approach addresses known limitations in computational protein design where individual metrics may be insufficient for predicting experimental success [32].

### 4.3 Structural Biology and Biophysical Insights

The binding affinity differences among analogues provide insights into epitope tolerance and functional constraints. The observation that BM2 and BM8 maintain highest affinities (77 and 88 nM) while achieving successful solubilization suggests optimal balancing of epitope preservation with solubility enhancement.

#### 4.3.1 Environmental and Stability Factors Influencing Binding Affinity

The superior binding performance of BM2 and BM8 relative to other analogues can be attributed to several interconnected factors beyond simple sequence conservation. First, the local chemical environment around the epitope regions differs significantly between analogues due to their distinct amino acid substitution patterns. BM2 and BM8 may have achieved more favorable electrostatic and hydrophobic environments that better support the native-like presentation of critical epitope residues, even in aqueous solution.

#### 4.3.2 Comparison with Native Membrane Environment

The difference between our measured K_D_ values (77-857 nM) and literature values (28.4 pM) for native mAb1-CCR8 binding reflects well-documented effects of membrane environment on GPCR binding properties [30, 31]. Studies consistently show that detergent solubilization can reduce apparent binding affinities by 10-1000 fold compared to native membrane presentation, due to altered protein dynamics, loss of membrane-mediated interactions, and conformational changes induced by detergent environments.

### 4.4 Therapeutic and Biotechnological Implications

The successful solubilization of CCR8 while preserving antibody binding opens new avenues for therapeutic development. Soluble analogues enable standard biochemical assays, high-throughput screening approaches, and structural biology studies that are challenging with membrane-embedded proteins. This accessibility could accelerate development of CCR8-targeted immunotherapies for cancer treatment and autoimmune disease management.

The demonstrated epitope preservation suggests potential applications in antibody discovery and optimization. Soluble analogues could serve as immunogens for antibody generation, screening targets for antibody libraries, and platforms for epitope mapping studies. The high solubility and preserved epitope presentation of these analogues make them particularly valuable for de novo binder generation approaches, including phage display, yeast display, and computational antibody design platforms, where the ability to work in aqueous conditions without detergents significantly simplifies screening workflows and improves hit identification rates. The high-yield production (up to 73.72 mg/L) makes large-scale applications feasible for industrial antibody development programs. Beyond traditional drug discovery applications, soluble CCR8 analogues could enable novel therapeutic strategies including high-throughput small molecule screening, allosteric modulator discovery, and functional assays in simplified biochemical systems that avoid the complexity and cost of cell-based approaches. Soluble receptor analogues could also function as competitive binding partners that modulate the tissue distribution and clearance kinetics of CCR8-targeted therapeutics, like strategies employed with soluble VEGFR decoys like aflibercept. Such approaches could potentially enhance drug delivery to tumor sites while reducing off-target effects in healthy tissues expressing CCR8 [35].

### 4.5 Platform Technology and Broader Applications

This work establishes a validated platform technology applicable to other membrane protein targets. The systematic approach combining computational design with comprehensive experimental validation provides a roadmap for solubilizing challenging membrane proteins while preserving functional properties. The methodology could be particularly valuable for other GPCRs, ion channels, and transporters where membrane environment currently limits structural and functional studies.

Recent advances in computational protein design, including RFdiffusion-based approaches and enhanced ProteinMPNN variants, suggest continued improvements in success rates and design quality [33, 34]. Integration of these emerging tools with the validated workflow presented here could further enhance membrane protein solubilization capabilities.

### 4.6 Limitations and Future Directions

Several limitations warrant consideration for future studies. The binding affinity reduction compared to native membrane presentation represents a trade-off inherent in solubilization approaches. Future work could explore membrane mimetic systems or lipid nanoparticle incorporation to maintain more native-like binding properties while preserving solubility benefits.

The exclusive focus on mAb1 binding, while validating epitope preservation, leaves broader functional characterization unexplored. Future studies should examine G-protein coupling, downstream signaling, and interactions with other ligands to comprehensively validate functional preservation. Additionally, structural studies using cryo-EM or X-ray crystallography would provide atomic-level validation of design accuracy and guide further optimization efforts.

Long-term stability, thermal tolerance, and aggregation propensity represent important practical considerations for biotechnological applications. Systematic characterization of biophysical properties could inform formulation strategies and guide selection of optimal analogues for specific applications.

### 4.7 Broader GPCR and Channel Protein Applications

The validated computational pipeline could be readily applied to other therapeutically relevant GPCRs including CCR5 (HIV entry), CXCR4 (cancer metastasis), and dopamine receptors (neurological disorders). The systematic approach of epitope preservation combined with solubilization could accelerate drug discovery timelines across the broader GPCR family, particularly for targets where structural information remains limited due to expression and stability challenges.

## 5. Conclusion

This work demonstrates a computational strategy for engineering a highly hydrophobic GPCR, CCR8, into soluble analogues that maintain an antibody-binding epitope. By integrating structural modeling, solubility optimization, and sequence filtering, we generated variants that express in E. coli, purify under aqueous conditions, and bind mAb1 on yeast surface display. Eight designs showed measurable binding, with the best-performing analogues achieving affinities comparable to wild-type CCR8 in detergent micelles. These results establish a practical framework for generating soluble GPCR variants as functional surrogates for antibody screening and characterization, reducing reliance on detergents or membrane mimetics. Future efforts will focus on structural validation and expanding design constraints to include ligand-binding functionality, enabling broader applications in GPCR-targeted drug discovery.

This study successfully demonstrates computational design and experimental validation of soluble CCR8 analogues that maintain native antibody binding properties. The 62% experimental success rate, combined with preserved binding affinities in the 77-857 nM range, validates the effectiveness of ProteinMPNN-sol sequence design for challenging membrane protein targets.

The extensive sequence divergence (10-13% identity) while maintaining functional epitope presentation demonstrates the robustness of computational constraint strategies and provides confidence in broader applicability to other membrane protein systems. The high-yield production of multiple analogues (1.19-73.72 mg/L) establishes practical feasibility for biochemical and therapeutic applications.

This work represents a significant advancement in membrane protein accessibility, addressing fundamental limitations that have historically hindered structural studies and therapeutic development. The validated methodology provides a platform technology for solubilizing other challenging membrane proteins while preserving critical functional properties, with immediate applications in CCR8-targeted immunotherapy development and broader implications for membrane protein drug discovery.

## Appendix

**Supplementary Figure 1.**
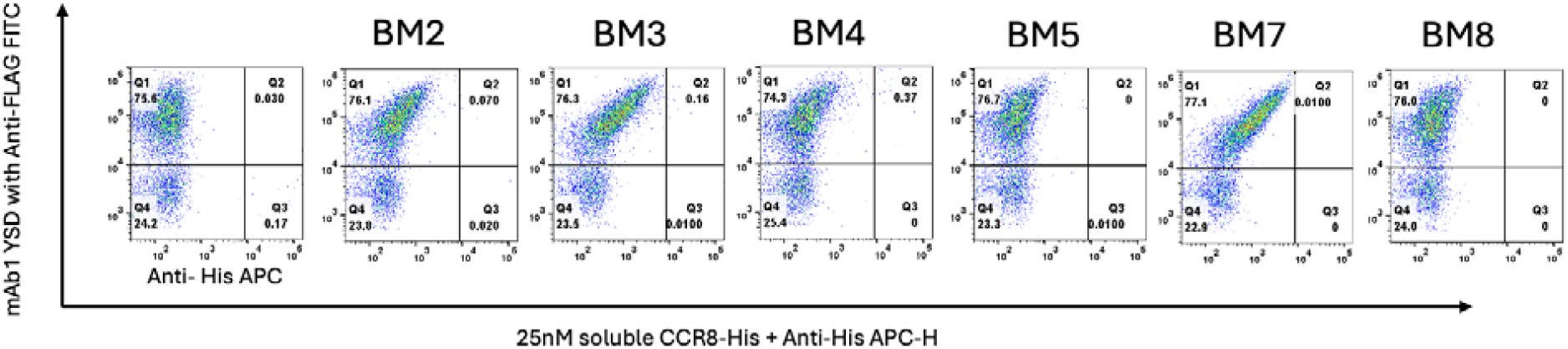
Flow Cytometry Analysis of CCR8 Analogues Binding at 25 nM mAb1 Concentration. Flow cytometry plots demonstrating the binding of 25 nM CCR8 analogues (BM2, BM3, BM4, BM5, BM7, BM8) and 50 nM WT CCR8 to surface-displayed mAb1 (scFv form). The y-axis shows the display efficiency of mAb1 on the yeast surface, while the x-axis represents mAb1 binding to His-tag CCR8 analogues detected via His tag antibody, Alexa Fluor 647. This lower concentration analysis complements the 250 nM data shown in Figure 3, demonstrating dose-dependent binding across the analogue series.

**Supplementary Figure 2.**
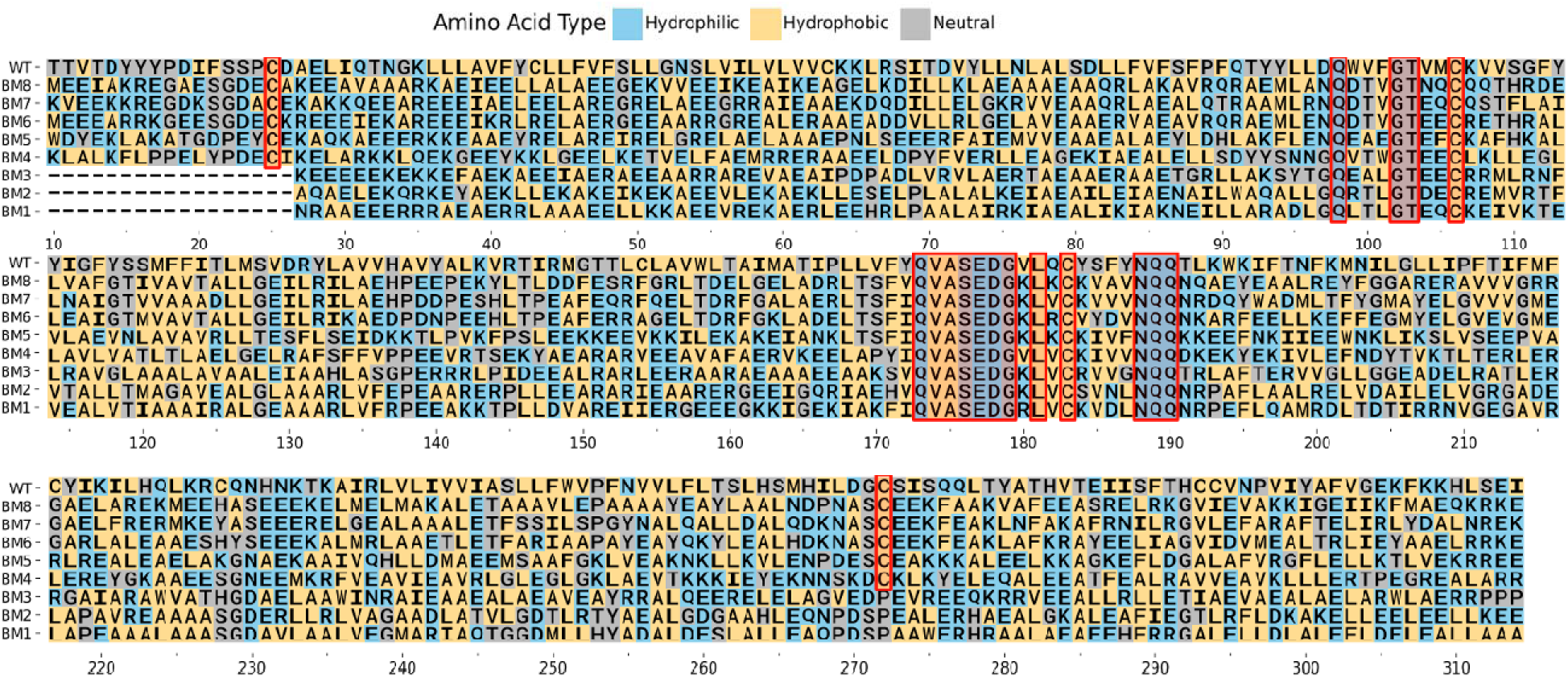
Sequence Alignments of CCR8 Soluble Designs with Wild-Type CCR8. Amino acids are colored based on the IMGT Hydropathy classes. The x-axis indicates the positions of wild-type CCR8 (start from 10), and fixed residues are highlighted with red box.

